# Generating human neural diversity with a multiplexed morphogen screen in organoids

**DOI:** 10.1101/2023.05.31.541819

**Authors:** Neal D. Amin, Kevin W. Kelley, Jin Hao, Yuki Miura, Genta Narazaki, Tommy Li, Patrick McQueen, Shravanti Kulkarni, Sergey Pavlov, Sergiu P. Paşca

## Abstract

Morphogens choreograph the generation of remarkable cellular diversity in the developing nervous system. Differentiation of stem cells toward particular neural cell fates *in vitro* often relies upon combinatorial modulation of these signaling pathways. However, the lack of a systematic approach to understand morphogen-directed differentiation has precluded the generation of many neural cell populations, and knowledge of the general principles of regional specification remain incomplete. Here, we developed an arrayed screen of 14 morphogen modulators in human neural organoids cultured for over 70 days. Leveraging advances in multiplexed RNA sequencing technology and annotated single cell references of the human fetal brain we discovered that this screening approach generated considerable regional and cell type diversity across the neural axis. By deconvoluting morphogen-cell type relationships, we extracted design principles of brain region specification, including critical morphogen timing windows and combinatorics yielding an array of neurons with distinct neurotransmitter identities. Tuning GABAergic neural subtype diversity unexpectedly led to the derivation of primate-specific interneurons. Taken together, this serves as a platform towards an *in vitro* morphogen atlas of human neural cell differentiation that will bring insights into human development, evolution, and disease.

## Introduction

Human stem cell biology for neuroscience offers the promise to generate a diversity of neural cell types of interest to study human development ^3-5^, identify the pathophysiology of neuropsychiatric disorders ^6-9^, and develop cell-based therapies ^10-12^. By mimicking *in vivo* morphogen signaling events *in vitro*, human pluripotent stem (hPS) cells can be guided to differentiate into specific neuronal and glial populations. For example, the application of sonic hedgehog (SHH), which is secreted by floor plate cells located in the ventral neural tube, guides stem cells to ventral neural cell fates ^13-18^. In fact, protocols combining SHH with WNT and FGF8 agonists yield human stem cell-derived midbrain dopaminergic neurons that have enabled recent transplantation clinical trials for Parkinson’s disease ^19,20^. Alternatively, combining SHH with WNT and retinoic acid (RA) generates human spinal motor neurons that have been used to model motor neuron disease ^21^.

Generally, these protocols have been developed by taking knowledge of spatiotemporal morphogen expression patterns and iteratively modulating morphogen pathways *in vitro* by trial and error to generate a neural population of interest. Unfortunately, our understanding of human morphogen signaling is incomplete, and it is still not clear how a limited set of signaling pathways give rise to the immense cell diversity in the nervous system. Furthermore, approaches to differentiate hPS cells have not been comprehensive, and even with the advent of guided and unguided organoid differentiation approaches ^22^, we have yet to generate many neural cells *in vitro*.

Towards this goal, we built an arrayed morphogen screening platform that leverages regionalization of human neural organoids, multiplexed single cell RNA sequencing and transcriptomic mapping onto reference atlases of the human developing nervous system. Using this approach, we found that discrete molecular conditions generated highly regionalized neural organoid cultures that collectively cover neural diversity across the neural axis. We developed and applied bioinformatic analyses to deconvolute morphogen-cell type relationships. This enabled us to extract principles of morphogenesis including discrete timing windows for patterning and a protocol to generate a rare human interneuron subtype recently identified as primate-specific ^24,25^. Ultimately, the promise of this scalable platform is to build a comprehensive morphogen atlas of human neural cell fate specification to accelerate human research into neurobiology, cellular pathophysiology, and cell-based therapies across nervous system disorders.

## Results

Morphogens expressed from neurodevelopmental organizer cells choreograph neural cell fate specification across spatial domains of the neural axis (Fig. 1A) ^2^. To understand the principles of morphogen action upon human neural differentiation, we designed a screen in human induced pluripotent stem (hiPS) cell-derived organoids (Fig. 1B). We first generated spheroids from hiPS cells in microwells and neuralized them with dual SMAD inhibitors, as previously described ^26-28^. We screened modulators of 8 morphogen pathways using 14 different molecules in 46 unique combinations, timings, concentrations, and durations in an arrayed format (Fig. 1C, **Fig. S1A, Table S1**). To facilitate the standardized application of morphogens, we designed and fabricated polylactic acid (PLA) polymer grids with 3D printing that separated 30 organoids per condition in 6-well plates, allowing us to grow a panel of nearly 1500 individually spaced organoids (**Fig. S1B**). We tracked organoid size over time, and at days 72-74 of differentiation, we used tissue clearing methods to examine organoid morphological features and dissociated organoids for multiplexed single cell RNA sequencing (scRNA-seq) using split-pool combinatorial barcoding ^29^.

**Figure 1.**
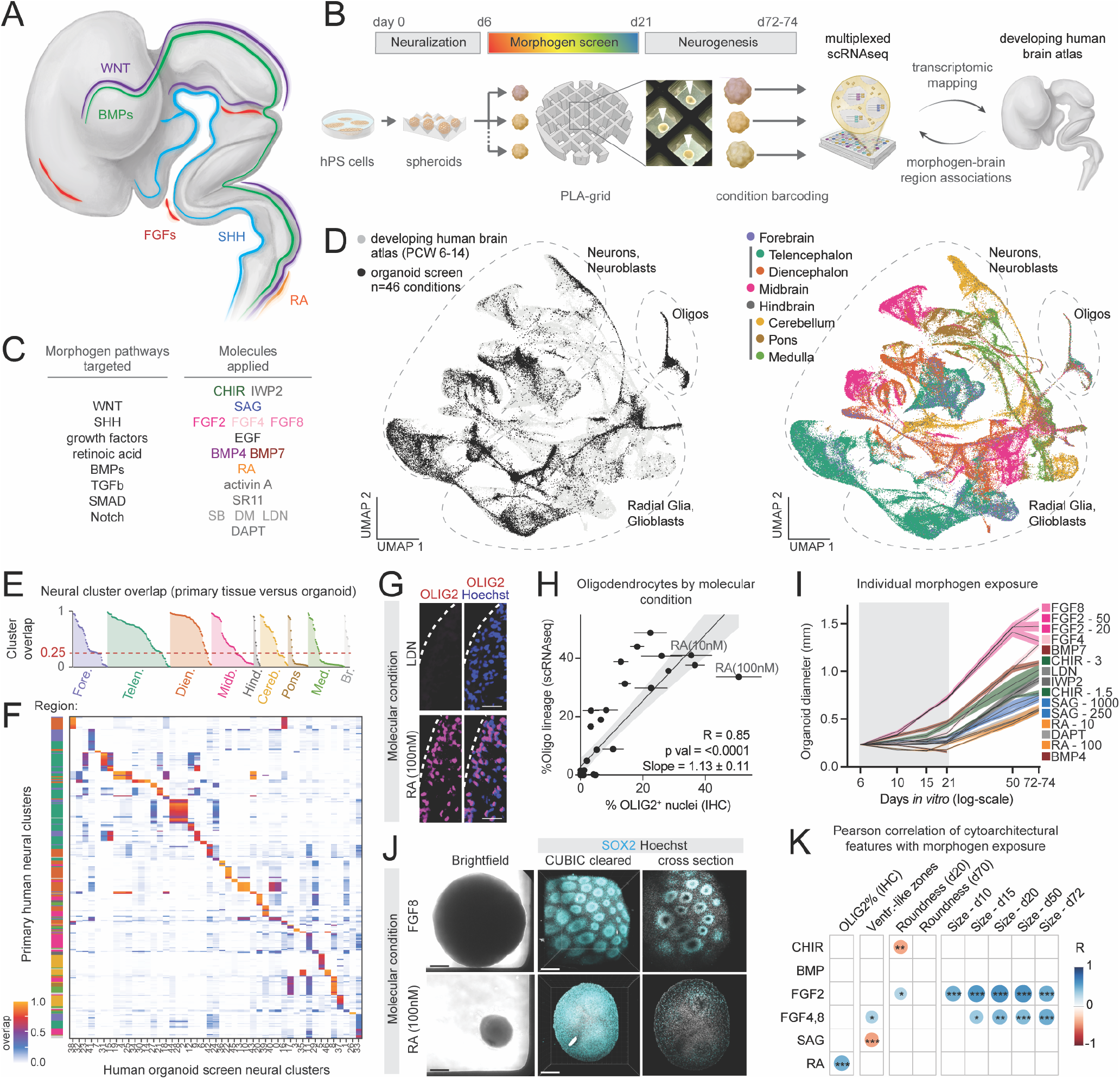
Generation of human neural diversity with a multiplexed morphogen screen in organoids. **(A)** Schematic of morphogen expression in the developing human fetal nervous system (adapted from 2). **(B)** Diagram of the arrayed screening platform. White arrows indicate human brain organoids within the polylactic acid (PLA) grid. **(C)** Molecules used to modulate morphogen pathways. SAG, Smoothened agonist; RA, retinoic acid; DM, dorsomorphin; SB, SB-431542; LDN, LDN-193189; SR11, SR11237; FGF, fibroblast growth factor; EGF, epidermal growth factor; SHH, sonic hedgehog; TGF, transforming growth factor; BMP, bone morphogenic protein; CHIR, CHIR99021. **(D)** UMAP of integrated cellular profiles from organoid morphogen screen and human fetal brain atlas 23 colored by dataset (left) and fetal brain region (right). **(E)** Overlap of co-clustered primary and organoid single cell clusters ordered by fetal brain region (1 indicates perfect cluster overlap in the integrated dataset). **(F)** Heatmap of cluster overlap between integrated primary and organoid cell clusters. Primary fetal clusters with overlap < 0.25 to organoid clusters are not shown. **(G)** Immunohistochemistry of OLIG2 expression in d72 organoids treated with LDN or 100nM retinoic acid (RA). (25 nm, scale bar). **(H)** Scatter plot of the percentage of OLIG2+ nuclei by IHC versus percentage of oligodendrocytes by scRNA-seq in organoids from each condition. (mean, standard error, linear regression, 95% confidence interval). **(I)** Mean organoid diameter from conditions exposed to a single molecule (standard error) over time. One condition treated with BMP4 alone decreased in size and disintegrated. **(J)** Brightfield and CUBIC-cleared organoids from FGF8 and 100nM RA-treated d72 organoids. (scale bar, 0.2mm). **(K)** Pearson correlation of quantitative morphological and cellular organoid features with the presence of each molecule across conditions. Only associations that pass P-value cut-offs are shown ^*^<0.05, ^**^<0.01, ^***^<0.001, Pearson correlation.

After filtering, we successfully captured 36,265 high quality cells across 46 organoid conditions (average of 788 ± 57 (standard error of the mean) cells per condition, **Fig. S1C**). To map cellular diversity, we integrated cells from all conditions with a recently published comprehensive developing human fetal brain transcriptomic atlas ^23^ and visualized cells in two-dimensional space using uniform manifold and approximation projection (UMAP) (**Fig. 1D**). Cells from the organoid screen spanned neuronal, astroglial, and oligodendrocyte cell types from the telencephalon to the hindbrain (**Fig. 1D, Fig. S1D**). In the integrated dataset, we examined similarity of single cell clusters from primary fetal brain data with clusters from the organoid screen (**Fig. 1E**). Overall, organoid clusters mapped to 65% of fetal central nervous system derived single cell populations with at least 25% overlap (**Fig. 1F**, Methods), indicating we captured a large proportion of brain regional diversity found *in vivo*.

To assess the validity of the multiplexed scRNA-seq data, we used immunohistochemistry (IHC) to stain for the transcription factor OLIG2, a marker of oligodendrocyte precursor cells (OPC) and some ventral neural tube progenitors ^30^ (**Fig. 1G**). Within each condition, we found high consistency of the fraction of OLIG2^+^ nuclei in cryo-sectioned organoids (**Fig. S2A,B**). Importantly, OLIG2 percentage by IHC and the percentage of oligodendrocyte lineage cells determined by scRNA-seq were correlated across conditions (r = 0.85, P< 0.0001, **Fig. 1H**). Conditions that contained RA and the SHH agonist SAG generated the highest numbers of oligodendrocyte lineage cells compared to conditions without these molecules (**Fig. S2B**), consistent with prior reports in organoids and 2D stem cell cultures ^31-33^. These results demonstrate that this platform can make quantitative assessments of morphogen effects on organoid cell composition.

In addition to affecting cell type composition, molecular signaling in organoids can influence cellular proliferation and morphology. Thus, we used imaging to extract organoid silhouette features over time. We found that organoid growth rate and shapes are related to morphogen exposure (**Fig. 1I, Fig. S2C-F, Table S2**). Among conditions containing a single morphogen modulator, application of FGF8 (100 ng/mL) resulted in organoids with an average diameter that was 2.82 times (P<0.0001) larger than those exposed to RA (100 nM). To assess organoid morphology differences, we also performed organoid clearing with CUBIC and whole mount SOX2 staining ^34^ (**Fig. S2H**). Strikingly, organoids exposed to FGF8 contained numerous ventricular-like regions compared with organoids exposed to other molecules such as RA (**Fig. 1J, Fig. S2G**). We further performed a multiple correlation analysis to identify associations between molecular pathway activation and organoid cytoarchitecture (**Fig. 1K**). We found that FGFs were positively associated with organoid size with FGF4 and FGF8 having the greatest positive correlation with the number of ventricular-like regions. OLIG2^+^ percentage was most positively correlated with the presence of RA. These results demonstrate that a comprehensive screening approach can extract physical and cell composition features associated with morphogen signaling in organoids.

We next examined scRNA-seq clusters to investigate cell type transcriptional identity in the arrayed multiplexed data. Using cell type marker gene expression, we discovered the generation of neuronal and non-neuronal cell populations spanning neurotransmitter subtypes and glial lineages (**Fig. 2A,B; Fig. S3A; Table S3**). Expression of canonical regional markers demonstrated that these cell types spanned forebrain, midbrain, and hindbrain regions (**Fig. 2C, Fig. S3B**). Transfer labeling from the reference primary human fetal regional annotations further corroborated diverse regional identity of organoid cells (**Fig. S3C-E**). This was also confirmed using VoxHunt ^35^ to map cell clusters to developing mouse brain *in situ* hybridization data, which identified organoid clusters corresponding to telencephalon, diencephalon, midbrain, and cerebellar regions (**Fig. S3F**). Finally, we integrated the results from each of these analyses to annotate the cell populations generated in the arrayed screen (**Fig. 2D, Table S3**). This revealed clusters of forebrain glutamatergic neurons and GABAergic neurons, as well as cerebellar Purkinje neurons and granule cells, *TP73*^*+*^*/RELN*^+^ Cajal-Retzius (CR) cells, and thalamic *VGLUT2*^+^ and corticotropin releasing hormone (CRH) expressing neurons (**Fig. S4A-B**). We also generated diverse populations of non-neuronal cell types such as *TTR*^*+*^*/CLIC2*^+^ choroid plexus cells, *MOG*^+^ maturing oligodendrocytes, and cortical hem cells (**Fig. S4A-C**).

**Figure 2.**
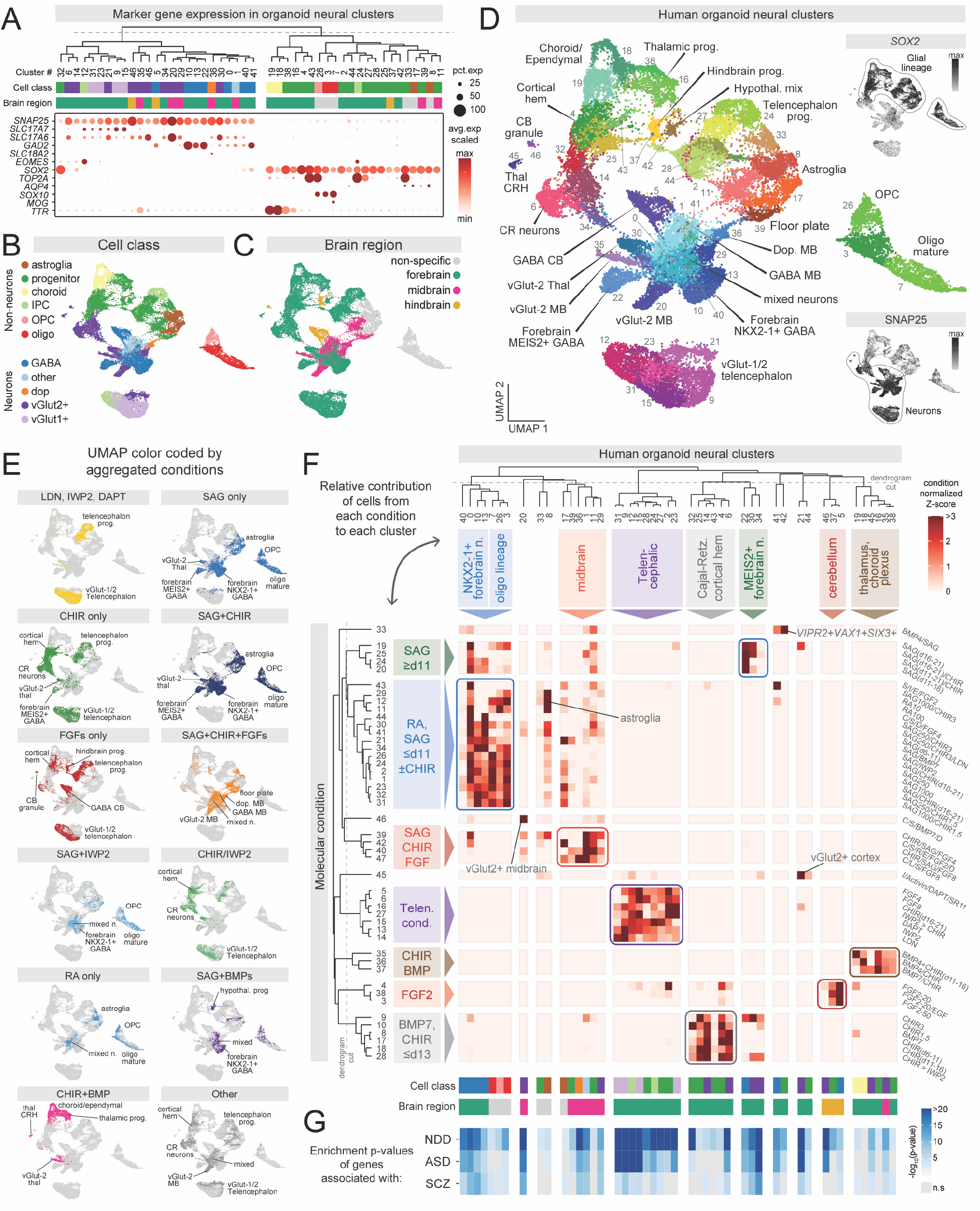
Deconvoluting morphogen modulators driving cellular composition. **(A)** Dot plot of cell class gene expression in single cell clusters from all organoid conditions. Top: hierarchical clustering based on top 20 cluster marker genes. **(B)** UMAP visualization of major cell class annotations. **(C)** UMAP visualization of brain regions annotations. **(D)** Annotated clusters of neural cells from all organoid conditions. Insets show gene expression of indicated markers. **(E)** UMAP visualization of aggregated molecular conditions sharing the indicated molecule(s). **(F)** Heatmap of cell composition (normalized by total number of cells in each condition) by molecular condition and scRNA-seq cluster. Groups of conditions that similarly contribute to sets of single cell clusters are color-coded and annotated. Dendrograms based on hierarchical clustering. **(G)** Enrichment p-values (hypergeometric test) of disease genes compared to cluster marker genes. Significance threshold determined from Bonferroni correction for the number of clusters.

We next inspected cell diversity generated from each molecular condition. Conditions exposed to high or low concentrations of the same molecule generated cells that overlapped in UMAP space (Fig. S5A). Conditions exposed to IWP2, LDN, or DAPT, inhibitors of the WNT, SMAD, and Notch pathways, respectively, almost exclusively generated telencephalic progenitors and glutamatergic neurons, while activation of SHH signaling resulted in forebrain GABAergic neurons, astrocytes, and oligodendrocytes (Fig. 2E). We quantified the relative generation of unique cell populations by aggregate molecular condition and found that certain populations such as dopaminergic neurons and choroid plexus were only generated when organoids were exposed to specific combinations of molecules, SAG+CHIR+FGF8 and BMP+CHIR, respectively (as expected from prior studies ^20,36-39^) (Fig. S5B). Progenitors, *VGLUT1*^+^ neurons, *VGLUT2*+ neurons, and GABAergic neurons were present more broadly across conditions with shared molecular features (Fig. S5C).

To examine associations between morphogen conditions and neural patterning at higher resolution, we quantified the contribution of cells from each condition to single cell clusters to identify condition-to-cell type associations (Fig. S5C, Table S4). On average, each condition contributed at least 10% of cells to 3.37 (± 0.16, standard error) neural clusters, reflecting the specificity of cell type generation within organoids from a single condition. Furthermore, most single cell clusters were populated with cells originating from more than one condition. By calculating a similarity matrix of conditions based upon contribution to clusters, we identified molecular conditions that generated organoids with similar cellular composition, allowing us to examine shared features of molecule identity, timing, concentration, and combination that led to neural cell formation (**Fig. 2F, Fig. S5D)**. Interestingly, we found groups of conditions that generated very similar neural populations including interneuron sub-types, midbrain, cerebellum, cortical hem, Cajal-Retzius cells, thalamus, and choroid plexus (Fig. 2F).

As stem cell-derived organoids are used to investigate the cellular phenotypes of genetic mutations associated with neuropsychiatric conditions ^40,41^, we examined the expression of genes with coding variation associated with neurodevelopmental disorders (NDD)^42^, autism spectrum disorder (ASD)^42^, and schizophrenia (SCZ)^43^ in cells generated in the organoid screen (**Fig. S6A**). Disease genes were broadly expressed and enriched across clusters with on average 81% (n= 373 genes), 84% (n= 184 genes), and 60% (n= 244 genes) of disease genes expressed per cluster from NDD, ASD, and SCZ, respectively (Fig. 2G). We used a specificity index to visualize disease-associated genes with restricted expression across clusters (**Fig. S6B,C**). While many neuropsychiatric genes are ubiquitously expressed across neuronal and non-neuronal cells, others are found in only a subset of clusters that correspond to specific molecular conditions (**Fig. S6C**). For example, *EBF3*, which is associated with a NDD syndrome characterized by autistic features and cerebellar hypoplasia among other symptoms^44^, is expressed in organoid cerebellum Purkinje, glutamatergic midbrain, and Cajal-Retzius neurons (**Fig. S6D**). Similarly, *GATA3*, which is associated with NDD, and *NR4A2*, which is associated with SCZ, demonstrated restricted expression in organoid mid-brain GABAergic and dopaminergic neuron clusters, respectively (Fig. S6D). Thus, we generated a diversity of neural organoids that can be used to investigate disease-associated gene-phenotype relationships in the relevant human neural cell populations.

Our association analysis identified three conditions that yielded high percentages of cerebellar neurons, and we investigated these cell clusters more closely by performing transcriptomic mapping onto a recent dataset from the human fetal cerebellum ^1^ (**Fig. 3A**). Three primary cerebellar clusters mapped with high specificity onto cells from the screen, including cerebellar Purkinje neurons expressing *SKOR2/FOXP2/TFAP2A/B, PAX2*+ cerebellar interneurons, and granule neurons expressing *RBFOX3, CBLN1, RELN*, and *PAX6* (Fig. 3B, Fig. S7A). By deconvoluting morphogen identity with brain region generation, we found that FGF2 in the presence of N2 supplement was the dominant feature of molecular conditions that yielded these cerebellar populations, similar to a prior report ^45^ (**Fig. 3C**). Indeed, cerebellar progenitors and neurons were generated with high efficiency in three conditions containing FGF2 alone or in combination with EGF, but not in conditions with FGF4 or FGF8 (**Fig. 3D**). By immunostaining, we confirmed SKOR2^+^/TFAP2A^+^ nuclei in organoids from these conditions in addition to populations of PAX6^+^ cells (**Fig. 3E,F, Fig. S7B**).

**Figure 3.**
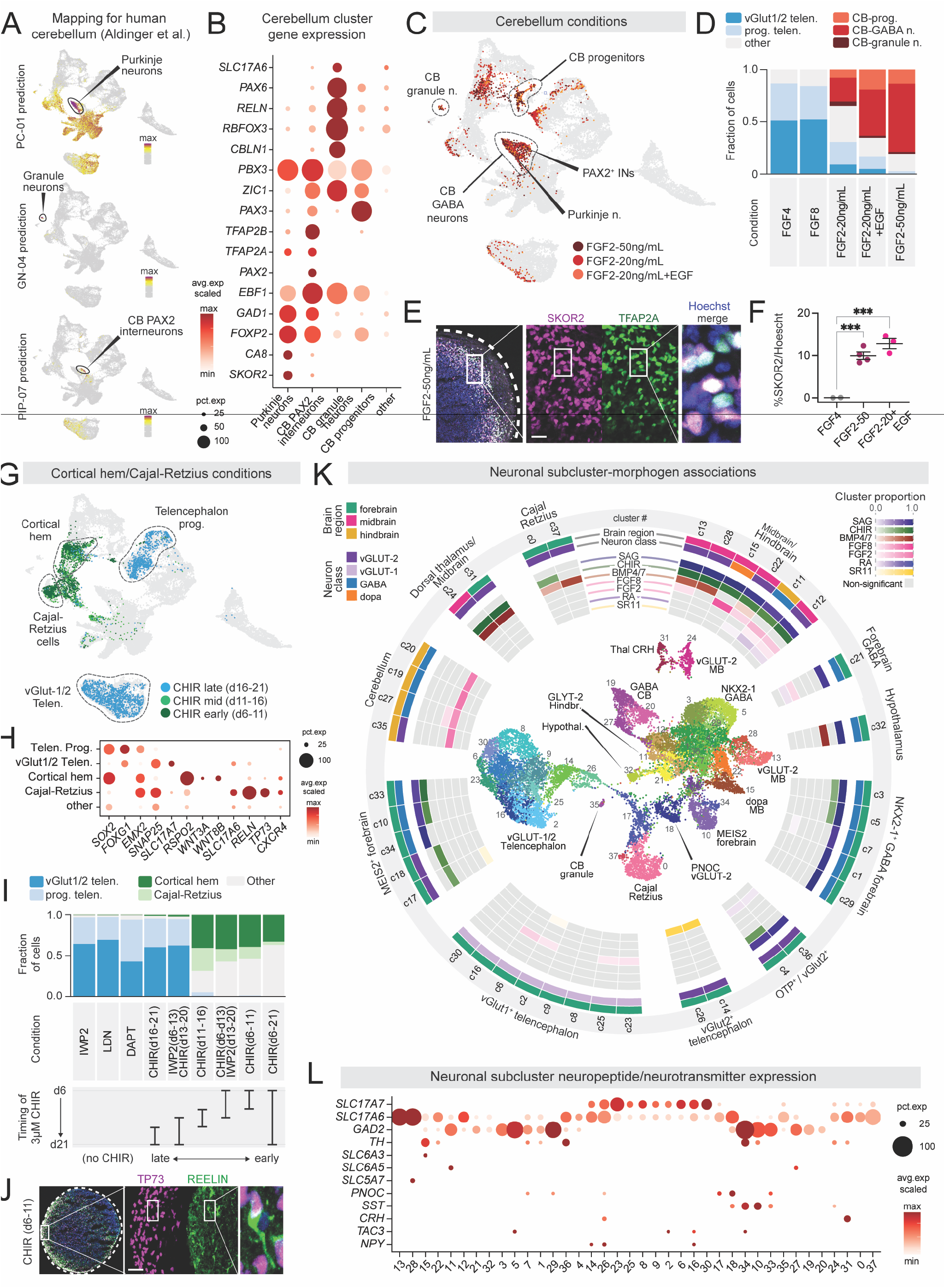
Extracting molecular features associated with unique cell types. **(A)** Label transfer of cerebellar populations ^1^ to scRNA-seq data identifies Purkinje cells, PAX2^+^ interneurons, and granule neurons. **(B)** Dot plot of marker gene expression in select clusters. **(C)** UMAP highlighting cells originating from three conditions exposed to FGF2 generating high numbers of cerebellar cells. **(D)** Quantification of cell proportions by scRNA-seq in the indicated conditions. **(E)** Immunohistochemistry for Purkinje cell transcription factor markers SKOR2 and TFAP2A. Scale bar, 25 nm. **(F)** Quantification of SKOR2 positivity in the indicated conditions. Each dot is a single organoid. ^*^<0.05, ^**^<0.01, ^***^<0.001 **(G)** UMAP highlighting cells from three conditions exposed to the WNT activator CHIR for 5 days at sequential time windows. **(H)** Dot plot of marker gene expression in select clusters. **(I)** Quantification of cell composition from scRNA-seq data, arranged by timing of CHIR application. **(J)** Immunohistochemistry of Cajal-Retzius cell transcription factor marker TP73 and cytoplasmic marker REELIN in organoid cryosection. Scale bar, 30 nm. **(K)** Outside: Neuronal scRNA-seq subclusters visualized in a circular heatmap of deconvoluted morphogen associations. Statistical associations based on permutation testing (methods). Inside: UMAP visualization of neuronal subclusters. **(L)** Dot plot expression of neurotransmitter associated genes and neuropeptides in each single neuronal subcluster.

When we inspected conditions that were exposed to the WNT activator CHIR, we observed that some conditions generated high percentages of glutamatergic cortical neurons, while others generated Cajal-Retzius (CR) neurons and cortical hem (CH), from which CR neurons principally derive (Fig. 3G,H, Fig. S7C)^46,47^. A closer examination of these molecular conditions revealed that CH/CR fates were specified when CHIR was applied starting early during differentiation, before day 11 (**Fig. 3I**). WNT activation from day 6 to 13 followed by WNT inhibition from day 13 to 21 still resulted in CR/CH cells, indicating that sustained WNT activation is not critical for this switch in cell fate. Interestingly, CHIR application starting after day 13 yielded glutamatergic cortical neurons, similar to conditions in which IWP2, LDN, or DAPT were applied. We confirmed the generation of CR cells by immunostaining which revealed TP73^+^ nuclei with cell bodies and processes expressing Reelin (**Fig. 3J**). Thus, this arrayed screen identified a surprisingly narrow critical timing window in which WNT activation can control distinct forebrain cell fates.

To examine neuronal cell type diversity in greater detail, we subclustered neuronal cells for further investigation. Using a permutation analysis, we found molecules statistically associated with the generation of each neuronal population, yielding a morphogen map of neuronal diversity (**Fig 3K, Table S5**). For example, thalamic neurons were generated from conditions in which CHIR with or without BMP7 or BMP4 were added (**Fig. 3K, Fig. S7E**). Furthermore, we identified a condition with the dorsalizing and ventralizing morphogens BMP4 and SAG that generated a population of cells enriched for *VIPR2, VAX1, FZD5*, and *SIX3* that are representative of the ventral hypothalamus (**Fig. S7D**). We performed differentiations of four stem cell lines to day 133 and observed expression of these and other hypothalamic markers by qPCR (Fig. S7F). Next, we identified the expression of a diverse array of neurotransmitter and neuropeptide-associated genes that mark distinct populations (**Fig. 3L, Fig. S8**). Interestingly, many genes are expressed in clusters with high specificity. For example, CRH expression was found in thalamic neurons in conditions applied with CHIR and BMP, and acetylcholinesterase (ACHE) expression was found in *ISL1*^+^/*PHOX2B*^+^ motor neurons from a condition in which SAG, CHIR and RA were applied (**Fig. S8B**). Thus, this arrayed screening platform generates great neuronal diversity, including some highly specialized neuronal populations.

We observed that our screen generated a large proportion of neurons expressing the GABA synthesizing enzyme and we subclustered these cells for closer examination (**Fig. 4A, Fig. S9A**). Conditions with SAG with or without CHIR generated forebrain *DLX1*^+^ interneurons that segregated into two groups marked by *MEIS2*^+^ and *NKX2-1*^+^ transcription factors (**Fig. S9B-D**). Early application of 250 nM SAG preferentially generated *NKX2-1*^+^ interneurons which are located more ventrally *in vivo*, while late application of 250 nM SAG generated more *MEIS2*^+^ interneurons, located more dorsally *in vivo* (Fig. 4B), confirming previous observations^48^. When SAG was applied at the intermediate time point day 11, a mixture of both subtypes was observed, reflecting a graded effect of SAG timing on interneuron subtype generation (**Fig. 4C, Fig. S9B-D**).

**Figure 4.**
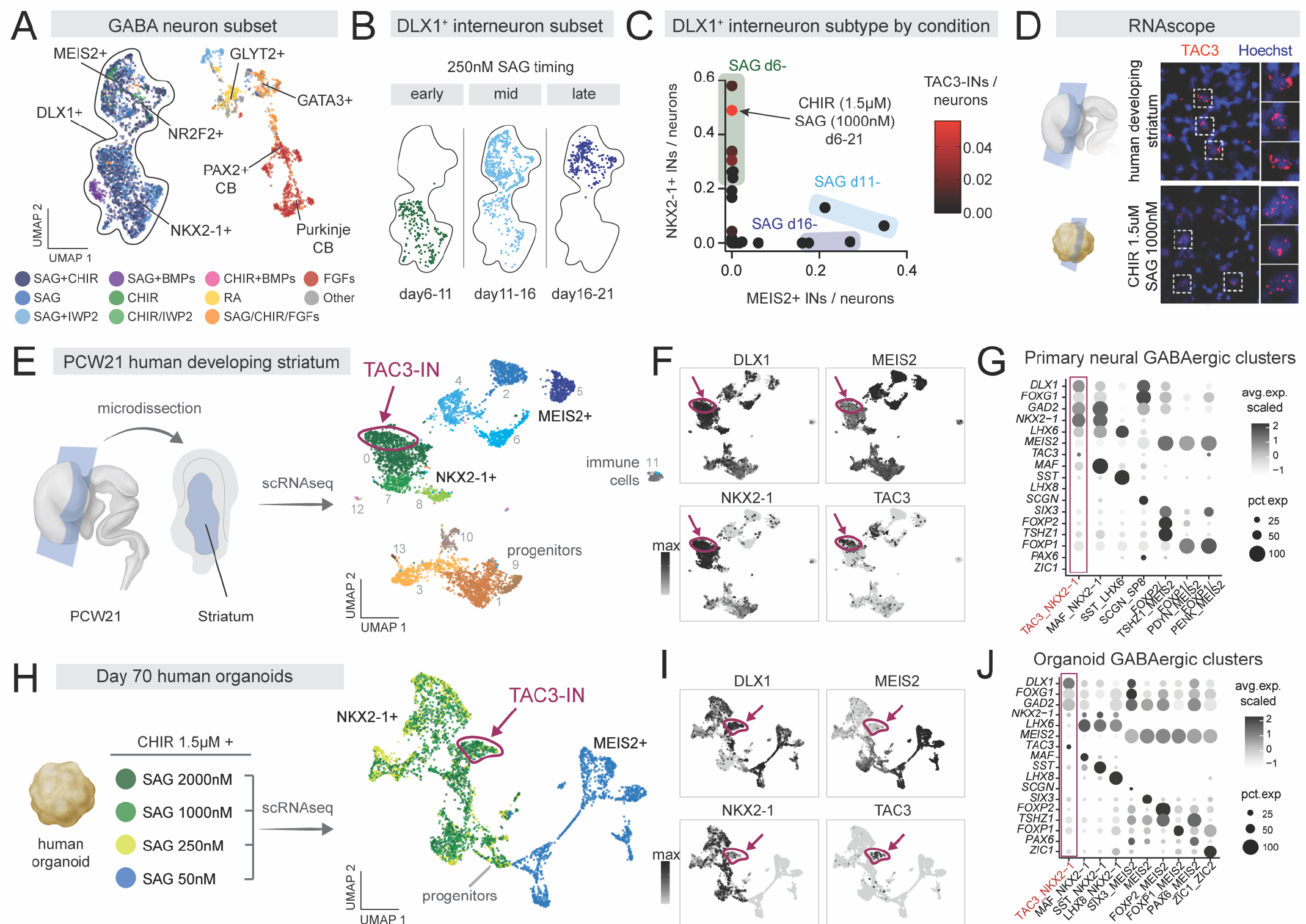
Conditions generating hPS cell-derived primate-specific TAC3^+^ interneurons. **(A)** UMAP of GABAergic neurons, color coded by aggregate molecular conditions and annotated by gene markers. **(B)** UMAP of DLX1^+^ forebrain interneuron clusters annotated by the timing of 250 nM SAG. **(C)** Scatter plot of percentage of MEIS2^+^ and NKX2-1^+^ interneurons generated in each condition. Conditions exposed to SAG early generate high percentages of NKX2-1+ interneurons and low percentage of MEIS2^+^ interneurons. Conditions are color coded by the percentage of TAC3-INs generated. **(D)** RNAscope of TAC3 in human primary striatal and organoid cells. **(E)** Left: diagram of PCW21 micro-dissected human striatum. Right: UMAP of scRNA-seq clusters highlights progenitors, GABAergic interneurons including TAC3-INs, and a small population of immune cells. Cells are color coded by cluster. **(F)** Interneuron marker gene expression (UMAP) in primary human cells with TAC3-INs highlighted. (G) Dot plot gene expression of previously identified TAC3 and striatal cell type marker genes in subpopulations of primary human fetal cells. **(H)** Left: diagram of organoid conditions exposed to 1.5 μM CHIR and 4 different concentrations of SAG. Right: UMAP visualization of day 73 scRNA-seq organoid conditions. Cells are color coded by condition. **(I)** Interneuron marker gene expression (UMAP) in human organoids with TAC3-INs highlighted. **(J)** Dot plot gene expression of previously identified TAC3 and striatal cell type marker genes in subpopulations of organoid cells.

When we examined neuropeptide diversity, we found tachykinin 3 (TAC3) was expressed in a subset of *NKX2-1*^+^/*DLX1*^+^ interneurons. Interestingly, a population of striatal-enriched TAC3^+^/DLX1^+^/NKX2-1^+^ forebrain interneurons (TAC3-INs) was recently identified in macaque and marmoset brains by single cell profiling ^24,25,49^. TAC3-INs were not found in striatum from mouse or ferret, suggesting this may be a primate-specific neuronal population. Of all conditions from the organoid screen, CHIR-1.5 μM and SAG 1000nM generated the greatest fraction of human TAC3-INs (Fig. 4C). We next prepared coronal sections of the developing human striatum from post conception week (PCW) 21 and performed RNAscope to confirm expression of TAC3 in primary and organoid cells (Fig. 4D). To further explore the transcriptomic identity of human TAC3-INs, we performed scRNA-seq on micro-dissected human primary striatum (**Fig. 4E, Table S6**). We identified a cluster of human *TAC3*^+^/*DLX1*^+^/*NKX2*-1^+^ neurons with gene expression patterns resembling those found in developing macaque and adult marmoset (Fig. 4F, G).

To further refine TAC3-IN generation from human stem cells, we performed new organoid differentiations in which we applied CHIR 1.5 μM and varied the concentration of SAG (Fig. 4H, Fig. S9E, Table S7). Low SAG (50 nM) generated *MEIS2*+ forebrain interneurons and higher SAG concentrations generated *NKX2-1*+ forebrain interneurons (Fig. 4I, Fig. S9F). The greatest fraction of TAC3-INs was observed with the highest concentration of SAG (2000 nM) (Fig. S9F). Previously reported marker genes for TAC3-IN from non-human primates showed similar expression patterns observed in TAC3-INs from hPS cell-derived organoids (**Fig. 4J, Fig. S9G**). Thus, we demonstrate that this organoid screening approach generated a reproducible protocol to further investigate a primate-specific human interneuron sub-type.

## Discussion

Advancing a mechanistic understanding of human cellular neurobiology and the pathophysiology of brain disorders relies upon building neuronal models that capture the cellular diversity of the human brain. Furthermore, development of cell therapies for conditions ranging from epilepsy to neurodegeneration requires detailed knowledge of how key brain cell types are generated from stem cells. To these ends, we bring together human brain organoid technology, multiplexed scRNA-seq, and transcriptomic mapping in an arrayed screen of morphogen modulators. This platform generated highly regionalized neural organoid cultures that collectively account for a large diversity of neuronal and non-neuronal brain cell types. We identified critical factors driving cerebellar differentiation and differential switch-like versus graded effects of morphogen timing on the generation of CR cells and GABAergic interneurons, respectively. Demonstrating the utility of this platform to identify and generate rare brain cell types, we characterized a new protocol yielding a recently described primate-specific forebrain interneuron subtype, TAC3-IN, that can be used to study evolutionary innovation in striatal circuits.

There are several limitations of this screen. There are multiple pathways controlling cell fate specification and the combinatorial possibilities to modulate them are immense. With the number of conditions included in this effort, we were able to generate a large, though incomplete, fraction of regional brain diversity. To build a comprehensive morphogen *vitro*, including activity-dependent processes that shape cell fate specification and maturation. Additionally, the effect of many morphogens will depend on pluripotency states and future studies should explore their impact across stem cell lines. Finally, comprehensive and well-annotated cellular atlases across human neurodevelopment will be needed to map cell types with higher precision.

We anticipate that technology developments, automation, and decreases in the cost of multiplexed sequencing over time will enable rapid scaling of this approach. Combining morphogens and transcription factor overexpression strategies may further accelerate efforts to generate cell states *in vitro* ^50,51^. Building larger datasets, mapping cells to reference atlases and applying advanced machine learning methods may ultimately lead to conditions that generate precise *in vitro* cell types in unexplored regions of the nervous system.

## Acknowledgements

This work was supported by the Stanford Brain Organogenesis Big Idea Grant from the Wu Tsai Neurosciences Institute (to S.P.P.), the NYSCF Robertson Stem Cell Investigator Award (to S.P.P.), the Kwan Research Fund (to S.P.P.), the Coates Foundation (to S.P.P.), the Senkut Research Funds (to S.P.P.), The Ludwig Foundation (to S.P.P.), the Chan Zuckerberg Initiative Ben Barres Investigator Award (to S.P.P.), CZ BioHub (to S.P.P.), NINDS K08-NS123544-01 (to N.D.A), Brain and Behavior Research Foundation (to N.D.A), Foundations of the National Institutes of Health Deeda Blair Research Initiative (to N.D.A.), NIMH T32MH019938 (to K.W.K), Stanford Maternal & Child Health Research Institute (MCHRI) Postdoctoral Fellowships (to Y.M.). We acknowledge A. Pasca and L. Li, (Stanford University) for assistance in processing tissue. This paper was typeset with the bioRxiv word template @Chrelli: www.github.com/chrelli/bioRxiv-word-template.

## Author contributions

N.D.A., K.W.K. and S.P.P. conceived the project and designed experiments. N.D.A and K.W.K performed experiments. J.H. assisted in organoid culture and performed RNAscope with K.W.K. and P.M. T.L. assisted in 3D printing. Y.M. obtained primary tissue single cell data. G.N. performed tissue clearing. S.K. assisted in qPCR and S.K. and S.P. assisted in image quantifications. N.D.A, K.W.K. and S.P.P. wrote the manuscript with input from all authors.

## Data and materials availability

Data is available at GEO accession number GSE233574.

## Materials and Methods

### Characterization and maintenance of hiPS cells

hiPS cell lines used were validated using methods as previously described. Genome-wide SNP genotyping was performed using Illumina genome-wide GSAMD-24v2-0 SNP microarray at the Children’s Hospital of Philadelphia (CHOP). Cultures were tested for and maintained Mycoplasma free. A total of 4 control hiPS cell lines derived from fibroblasts collected from 4 healthy subjects were used for experiments. Approval for this study was obtained from the Stanford IRB panel and informed consent was obtained from all subjects.

### Human primary tissue

Human brain samples were obtained under a protocol approved by the Research Compliance Office at Stanford University. The PCW21 tissue was dissected prior to dissociation for the scRNA-seq and RNA scope experiments.

### Organoid screen culture

Prior to neural differentiation, hiPS cells were cultured on vitronectin-coated plates (5 μg ml^−1^, Thermo Fisher Scientific, A14700) in Essential 8 medium with supplement (Thermo Fisher Scientific, A1517001). Cells were dissociated every 5 to 6 days with UltraPure™ 0.5 mM EDTA, pH 8.0 (Thermo Fisher Scientific, 15575) and passaged onto new plates. For the generation of 3D neural spheroids (day –1), hiPS cells were washed with PBS and incubated with Accutase (Innovative Cell Technologies, AT104) at 37°C for 7 min. Cells were manually triturated with a P1000 into a single cell suspension and were passed through a 40 μM filter (FloMi). To generate spheroids, 3 x 10^6^ hiPS cells were added to each AggreWell-800 well in Essential 8 medium supplemented with the ROCK inhibitor Y27632 (10 μM, Selleckchem, S1049), centrifuged at 100 *g* for 1 min, and then incubated at 37°C in 5 % CO_2_. After 24 hours, spheroids of approximately 10,000 cells were formed. Spheroids were lifted from each microwell by pipetting medium in the well up and down with a cut P1000 pipet tip and were placed in Essential 6 medium (Thermo Fisher Scientific, A1516401) with the SMAD pathway inhibitors dorsomorphin (2.5 μM, Sigma-Aldrich, P5499) and SB-431542 (10 μM, R&D Systems, 1614).

Approximately 200-300 spheroids collected from one AggreWell well were spread across 6 wells of an AggreWell plate, such that each well contained approximately 30 spheroids. Due to the microwell patterned bottom surface of the AggreWell, spheres within the same well did not come in contact. Between day 15 and day 21, growing organoids from a single AggreWell well were transferred to an ultra-low attachment 6-well plate (Corning, 3471) with a custom 3D fabricated polylactic acid grid inserted and adhered with inert silicone adhesive. Each organoid was placed within a single chamber of the grid. In brief, printing template was designed with Tinker-Cad (Autodesk 2023) and exported in a stereolithographic (stl) file. The printing parameters were as follows: layer height = 0.2 mm, minimum shell thickness (perimeter) = 0.6 mm (top) 0.7mm (bottom), nozzle temperature = 210C, bed temperature = 60C, and print speed = 75%. Grids were printed on a 3D printer (Prusa i3 MK3S+) using polylactic acid filament (Prusament by PLA White, PRM-PLA-JET-1000). After printing, grids were sterilized by soaking in 70% ethyl alcohol (Sigma-Aldrich, E7023) for one hour. Grids were then placed in a 2% solution of poly(2-hydroxyethyl methacrylate) (Sigma-Aldrich, P3932-10G) in ethanol to create a hydrophobic coating. Grids were allowed to dry in a sterile laminar flow hood and were UV treated for 30 minutes on each side. Grids were then inserted into each well of an ultra-low attachment 6-well plate (Corning, 3471) and a small amount (∼1g) of silicone adhesive (RS Hughes Inc RTV108, NC0380109) was applied around all sides to secure it to the plastic well. The silicone was allowed to cure for 24 hours. Wells with the 3D printed grids were washed 3 times in sterile PBS (Thermo Fisher Scientific, 14190250) prior to use.

Molecules in the screen were added to the following basal media: from day 0 to day 6, spheres were cultured in Essential 6 medium and media was exchanged daily. From day 6 to day 21, spheres were cultured in neural medium consisting of Neurobasal™-A Medium (Thermo Fisher Scientific, 10888022), B-27™ Supplement, minus vitamin A (Thermo Fisher Scientific, 12587010), GlutaMAX™ Supplement (1:100, Thermo Fisher Scientific, 35050079), and Penicillin-Streptomycin (1:100, Thermo Fisher Scientific, 15070063) and media was exchanged daily. From day 21 onwards, spheres were cultured in Neurobasal™-A Medium (Thermo Fisher Scientific, 10888022), B-27™ Supplement, minus vitamin A (Thermo Fisher Scientific, 12587010), GlutaMAX™ Supplement (1:100, Thermo Fisher Scientific, 35050079), N-2 supplement (Life Technologies, 17502048) and Penicillin-Streptomycin (1:100, Thermo Fisher Scientific, 15070063) with BDNF (20 ng/ml; Peprotech), NT3 (20 ng/ml; Peprotech), (cAMP; 50 μM, Millipore Sigma, D0627), L-Ascorbic Acid 2-phosphate Trisodium Salt (AA; 200 μM, Wako, 323-44822) and media was exchanged every 2-3 days.

Molecules used: EGF (R&D Systems, 236-EG), FGF2 (R&D Systems, 233-FB), IWP-2 (STEMCELL Technologies, 72124), RA (Sigma, R2625), 2.5 μM DAPT (STEMCELL Technologies, 72082), brain-derived neurotrophic factor (BDNF; 20ng ml-1, PeproTech, 450-02), NT3 (20 ng ml-1, PeproTech, 450-03), FGF4 (PeproTech, 100-31), FGF8 (PeproTech, 100-25), SAG (Millipore, 5666660), CHIR 099021 (Selleckchem, S1263), BMP4 (PeproTech, 120-05ET), BMP7 (PeproTech, 120-03P), LDN-193189 (S7507, Selleckchem), Activin A (50 ng/mL, PeproTech, 120-14P), SR11237 (100 nM, Tocris, 3411), dorsomorphin (DM, Sigma Aldrich P5499), SB (Tocris, 3748), Y-27643 (Selleckchem), L-ascorbic acid (FUJIFILM Wako Chemical Corporation, 344-44822), cAMP (Millipore Sigma, D0627).

### Cryoprotection and immunocytochemistry

Organoids were fixed in 4% paraformaldehyde (PFA)/phosphate buffered saline (PBS) overnight at 4°C. Organoids were then washed in PBS and transferred to 30% sucrose/PBS overnight. Organoids were rinsed in optimal cutting temperature (OCT) compound (Tissue-Tek OCT Compound 4583, Sakura Finetek) and frozen using dry ice. For immunofluorescence staining, 25 μm-thick sections were cut using a Leica Cryostat (Leica, CM1860). Cryosections were washed with PBS and blocked in 1% BSA, 0.1% Triton-X, sodium azide diluted in PBS for 1 h at room temperature. The sections were then incubated overnight at 4°C with primary antibodies diluted in 1% BSA, 0.1% Triton-X, sodium azide. Cryosections were washed three times and were incubated with secondary antibodies in 1% BSA, 0.1% Triton-X, sodium azide. Aquamount (Polysciences, 18606) was used to mount coverglass slides. Imaging was performed on a Leica TCS SP8 confocal microscope or Zeiss upright fluorescent microscope. Images were processed and analyzed using Fiji (NIH) and IMARIS (Oxford Instruments). To quantify percentage of oligodendrocyte precursor cells, images of Hoechst and OLIG2+ nuclei were taken. Hoechst images were converted into 8-bit images, performed automatic image threshold (Default, Dark Background) into a binary image, converted to Mask, performed Watershed (FiJi, NIH). OLIG2 nuclei images were manually thresholded due to inconsistency of background signals by a researcher blinded to molecular condition. Nuclei from Hoechst and OLIG2 images were quantified using Analyze Particles with pixel size 30-1000, and the number of OLIG2 nuclei counts were divided by Hoechst nuclei counts to obtain a percentage. Ventricular-like regions were quantified by examining cross sections of organoids stained with Hoechst and counting clearly identifiable ventricle-like structures.

Antibodies used: anti-reelin CR-50 (MBL International, D223-3), anti-P73 (EP436Y, Abcam, ab40658), Olig2 (AB9610, EMD Millipore), SKOR2 (NBP2-14565, Novus), SOX2 (D6D9, Cell Signaling Technologies), PAX6 (Invitrogen, MA1-109 (13B10-1A10)), MAP2 (Sigma, M4403-50uL), TFAP2A (sc-12726, Santa Cruz Biotechnologies), CLIC2 (EPR6469, ab175230, Abcam).

### Clearing and 3D staining of organoids

We applied the hydrophilic chemical cocktail-based CUBIC protocol. Organoids at day 72 were fixed with a 4% PFA/PBS solution at 4°C overnight, washed twice with PBS, and incubated in Tissue-Clearing Reagent CUBIC-L (TCI, T3740) at 37°C for 2 days. After washing three times in PBS, nuclei were stained with Hoechst. Samples were washed twice with PBS, and once with solution containing 10 mM HEPES, 10% TritonX-100, 200 mM NaCl and 0.5% BSA (HEPES-TSB) at 37°C for 2 hours, and then stained with anti-SOX2 (rabbit, Cell Signaling Technology, #3579, 1:100 dilution) antibody in HEPES-TSB solution at 37°C for 2 days. Samples were washed twice with 10% Triton X-100 in PBS and once with HEPES-TSB solution for 2 h each and then incubated with a donkey anti-rabbit IgG (H&L) highly cross-adsorbed secondary antibody, Alexa Fluor 488 (Thermo Fisher Scientific, A-21206, 1:300 dilution) in HEPES-TSB solution at 37°C for 2 days. Samples were washed twice with 10% Triton X-100 in PBS for 30 min and once with PBS for 1 hour. After washing with PBS, samples were incubated with Tissue-Clearing Reagent CUBIC-R+ (TCI, T3741) at room temperature for 2 d for refractive index matching. CUBIC-cleared organoids were then transferred into a well of a Corning 96-well microplate (Corning, 4580) in 150 μL of CUBIC-R+ solution and imaged using a 10x objective on a Leica TCS SP8 confocal microscope.

### Organoid morphology quantification

Brightfield images of individual organoids were taken at each time point from each condition. ROIs for all spheres were created manually using the polygon tool in ImageJ version 2.3.0/1.53f. For each resulting ROI, morphology features were analyzed using ImageJ. The resulting spreadsheets were merged into final data files with Python 3.8 using packages Pandas 1.5.2 and NumPy 1.23.5, from which figures were plotted in GraphPad Prism 9. Pearson correlations were performed with R studio package “corrplot”. Dendrograms were generated with Morpheus (Broad Institute, https://software.broadinstitute.org/morpheus), Sankey plots were generated with Flourish (Kiln Enterprises Ltd, UK).

### Real-time qPCR

mRNA from organoids were isolated using the RNeasy Mini kit (Qiagen, 74106) with DNase I, Amplification Grade (Thermo Fisher Scientific, 18068-015). Template cDNA was prepared by reverse transcription using the SuperScript™ III First-Strand Synthesis SuperMix for qRT-PCR (Thermo Fisher Scientific, 11752250). qPCR was performed using the SYBR™ Green PCR Master Mix (Thermo Fisher Scientific, 4312704) on a QuantStudio™ 6 Flex Real-Time PCR System (Thermo Fisher Scientific, 4485689).

### Single cell RNA-seq library preparation and data analysis

Three organoids from each condition were pooled at day 72–74, incubated in 30 U/mL papain enzyme solution (Worthington Biochemical, LS003126) and 0.4% DNase (12,500 U/mL, Worthington Biochemical, LS2007) at 37 °C for 45 minutes. Organoids were triturated with a P1000 pipette and were resuspended in FBS followed by trituration with a P200 pipette tip. Cells were resuspended in PBS with 0.4% DNase and filtered through a 70 μm Flowmi Cell Strainer (Bel-Art, H13680-0070). The samples were centrifuged at 200 x g for 3 minutes and were resuspended in Parse Cell Buffer containing BSA. Cells were then fixed using the Parse Biosciences Cell Fixation (Parse Biosciences, ECF2001) kit and stored at -80°C until the start of bar-coding and library prep with the Evercode Whole Transcriptome kit (Parse Biosciences, ECW02030). Samples were prepared with v1 kits, with the exception of SAG concentration series which was prepared with v2 kits. Libraries from different samples were pooled and sequenced by Admera Health on a NovaSeq S4 (Illumina).

To obtain single cell suspension from micro-dissected developing striatum, the tissue was incubated in 30 U/mL papain enzyme solution (Worthington Biochemical, LS003126) and 0.4% DNase (12,500 U/mL, Worthington Biochemical, LS2007) at 37 °C for 45 minutes. Then, tissue was washed with a solution including protease inhibitor after enzymatic dissociation, and gently triturated to obtain a single-cell suspension. Single cells were resuspended in 0.04% BSA/PBS (Millipore-Sigma, B6917-25MG) and filtered through a 70 μm Flowmi Cell Strainer (Bel-Art, H13680-0070), and the cell number was counted. To target 7,000 cells after recovery, approximately 11,600 cells were loaded per lane on a Chromium Single Cell 3′chip (Chromium Next GEM Chip G Single Cell Kit, 10x Genomics, PN-1000127) and cDNA libraries were generated with the Chromium Next GEM Single Cell 3’ GEM, Library & Gel Bead Kit v3.1 (10x Genomics, PN-1000128), according to the manufacturer’s instructions. The library was sequenced using the Illumina NovaSeq S4 2 × 150 bp at Admera Health.

### Single cell expression analysis

#### Preprocessing and quality filtering

For the multiplex organoid scRNA-seq screen, gene expression levels were quantified for each putative cell using the Parse Biosciences analysis software suite (split-pipe command, version 0.9.3p). Specifically, reads were mapped to a human (GRCh38, Ensemble release 93) reference genome created using the mkref mode and quantified using default parameters. All subsequent analyses were performed on the filtered count matrices using the R (version 4.1.2) package Seurat (version 4.3.0)(*52*).

To ensure only high-quality cells were included for downstream analyses, an iterative filtering process was implemented. First, cells with less than the 5^th^ percentile within the dataset of unique genes detected (785 genes) and with mitochondrial counts greater than 20% of the total counts were identified and removed. Subsequently, raw gene count matrices were normalized by regularized negative binomial regression using the sctrans-form function (vst.flavor=”v2”), which also identified the top 3,000 highly variable genes using default parameters. Dimensionality reduction using principal component analysis (PCA) on the top variable genes was performed and clusters of cells were identified in PCA space by shared nearest-neighbor graph construction and modularity detection implemented by the FindNeighbors and FindClusters functions using a dataset dimension of 50 (dims=50 chosen based on visual inspection of elbow plot) with default parameters. We subsequently performed iterative rounds of clustering (resolution = 2) to identify and remove clusters of putative low-quality cells based on outlier low gene counts (median below the 10^th^ percentile), outlier high fraction mitochondrial genes (median above the 95th percentile), outlier high fraction ribosomal genes (median above the 95th percentile), and/or high proportions of putative doublets identified by the Doublet-Finder package(*53*) (median DoubletFinder score above the 95th percentile). We iteratively removed low-quality clusters until no further clusters met the above thresholds. Following low quality cell removal, an additional round of filtering of putative stressed cells was performed using the Gruffi R package(*54*) in addition to manual inspection and removal of clusters expressing markers associated with unfolded protein response (e.g. *DDIT3, EIF2AK3*).

Analogous preprocessing and filtering were performed on the primary human fetal striatum and the secondary GABAergic neuron organoid screen scRNA-seq datasets, with the following exceptions. Filtered count matrices were obtained using the CellRanger analysis software suite (version 6.1.2) for the human fetal striatum dataset. Filtered count matrices were obtained using updated Parse Biosciences software for the GABAergic secondary screen (split-pipe command, 1.0.4p). Dataset dimension of 30 was chosen with cluster resolution of 1.

#### Cell clustering and annotation

Following low quality cell removal, datasets were clustered (FindClusters function; resolution = 2 for primary screen; resolution = 1 for primary fetal striatum and secondary GABAergic organoid screen) and embedded for visualization purposes with Uniform Manifold Approximation and Projection (UMAP)(*55*). Marker genes for each cluster were determined using the FindMarkers function with default parameters (Wilcoxon Rank Sum test), calculated on the normalized gene expression data. We identified and categorized major cell classes through a combination of marker gene expression and annotation via integration with a comprehensive reference human fetal brain datasets(*28, 44*). Specifically, progenitor clusters displayed expression of mitotic transcripts (e.g., *MKI67* and *TOP2A)* and/or *SOX2*, and had high cluster overlap to radial glial clusters annotated in fetal brain. Intermediate progenitor cells (IPC) expressed *EOMES* and mapped to fetal IPCs. We used the term astroglia to encompass multiple states of astrocyte differentiation and clusters expressed high levels of *SLC1A3* and *AQP4* and demonstrated mapping to fetal glioblasts. Oligodendrocyte progenitor cells (OPCs) expressed *PDGFRA* and *SOX10*, while oligodendrocytes expressed markers of myelination (*MOG, MYRF*). Glutamatergic neurons were identified by the presence of neuronal transcripts (*STMN2, SNAP25*) and presence of either VGLUT1 (*SLC17A7*) or VGLUT2 (*SLC17A6*). GABAergic neurons expressed GABA processing enzymes (GAD1, *GAD2*) and dopaminergic neurons expressed the dopamine transporter *SLC6A3*. Choroid-like cells were defined by expression of *TTR*. Similarly broad developmental region annotations were performed using a combination of integration/label transfer with the human fetal brain dataset and expression of known marker gene expression. Forebrain clusters were identified by expression of *FOXG1* (telencephalon), *EMX2, DLX1*, and/or *TCF7L2* (diencephalon) in addition to mapping to fetal forebrain, telencephalon, and diencephalon clusters. Mid-brain clusters expressed *EN1, OTX2, GATA3*, and/or markers of dopaminergic neurons in addition to mapping to fetal midbrain clusters. Hindbrain clusters primarily mapped to fetal hindbrain and cerebellum and contained markers of cerebellar cell populations.

To map our organoid clusters to annotated cell clusters from reference primary fetal single scRNA-seq data, we performed a pairwise dataset integration approach. Given the complex batch structure described in the Braun et al. fetal reference publication, we focused only on samples acquired using the 10x v3 chemistry. We further removed cell clusters annotated to non-central nervous system derived populations (erythrocyte, fibroblast, immune, neural crest, and placode annotated clusters) and removed cells from the earliest developmental stages which did not contain detailed regional information (samples from 6 PCW or younger). We used Seurat’s LogNormalize method (as implemented with the NormalizeData function) and integrated (IntegrateData function) using each donor as a batch (using dataset dimension of 50 and cluster resolution=1). The reference dataset was randomly subsetted to have a maximum of 500 cells per original cluster to improve computational efficiency. Using an analogous approach described previously (*56*), cluster overlap was defined as the proportion of cells in each integrated cluster that overlapped with the reference cluster labels. For the fetal cerebellum reference dataset, we downloaded processed data from UCSC cell browser, which was processed in the original study using similar workflow in Seurat. To further classify organoid cells, we utilized Seurat’s TransferData workflow to map reference regional annotation or fetal cerebellum clusters to our organoid cells.

For the primary fetal and organoid GABAergic secondary screen, annotation of clusters was based on marker genes and cell annotations used in a recent fetal macaque inhibitory neuron reference atlas (*22*).

#### Cell condition associations

To determine conditions associated with cell populations across the primary organoid screen clusters, we normalized the contribution of each condition to each cluster by the total number of cells present in each condition. To visualize positive associations, we scaled (to achieve mean = 0, standard deviation= 1) condition contributions to each cluster for each condition (visualized in Fig. 2F). For neuronal subclusters, we performed a permutation approach to calculate the statistical association of each applied molecule to a given neuronal subcluster. Specifically, for each cluster we calculated the proportion of cells that were exposed to a given molecule. To determine statistical associations for each molecule, we randomly shuffled molecular annotations (n= 1000 permutations) and determined p-values by calculating the number of permutated molecular proportions greater than the observed value and dividing by the number of permutations.

#### Disease gene enrichment

Neurodevelopmental disorder (NDD), autism spectrum disorder (ASD), and schizophrenia (SCZ) associated gene lists were compiled from recent rare coding variant association studies (*41, 42*). Using cutoffs defined by Fu et al., we focused on larger disease gene lists with the following criteria for each gene set: NDD (genes associated at FDR ≤ 0.001, 373 genes), ASD (genes associated at FDR ≤ 0.05, 185 genes), SCZ (genes associated at *P* < 0.01, 244 genes). To calculate enrichment of disease genes across organoid clusters, a one-sided Fisher’s exact test was performed for each set of cluster marker genes (adjusted P-value < 0.05) and disease gene sets. Focusing on genes with at least some minimal expression in our organoid data (defined as non-zero expression in at least 10% of cells in at least one organoid cluster) we determined which genes displayed the most specific expression across organoid clusters by calculating a Tau score. Tau varies from 0 to 1, where 0 means broadly expressed, and 1 is specific.

### Statistics

Data are presented as mean ± s.e.m. Raw data were tested for normality of distribution, and statistical analyses were performed by paired and unpaired t-test (two-tailed), Mann-Whitney test, Wilcoxon test, one-way *ANOVA*, or Kruskal-Wallis test with multiple comparison tests depend on the dataset. Sample sizes were estimated empirically. GraphPad Prism Version 9.3.1 or MATLAB (version R2018a, 9.4.0, MathWorks) were used for statistical analyses.

**Supplementary Figure 1.**
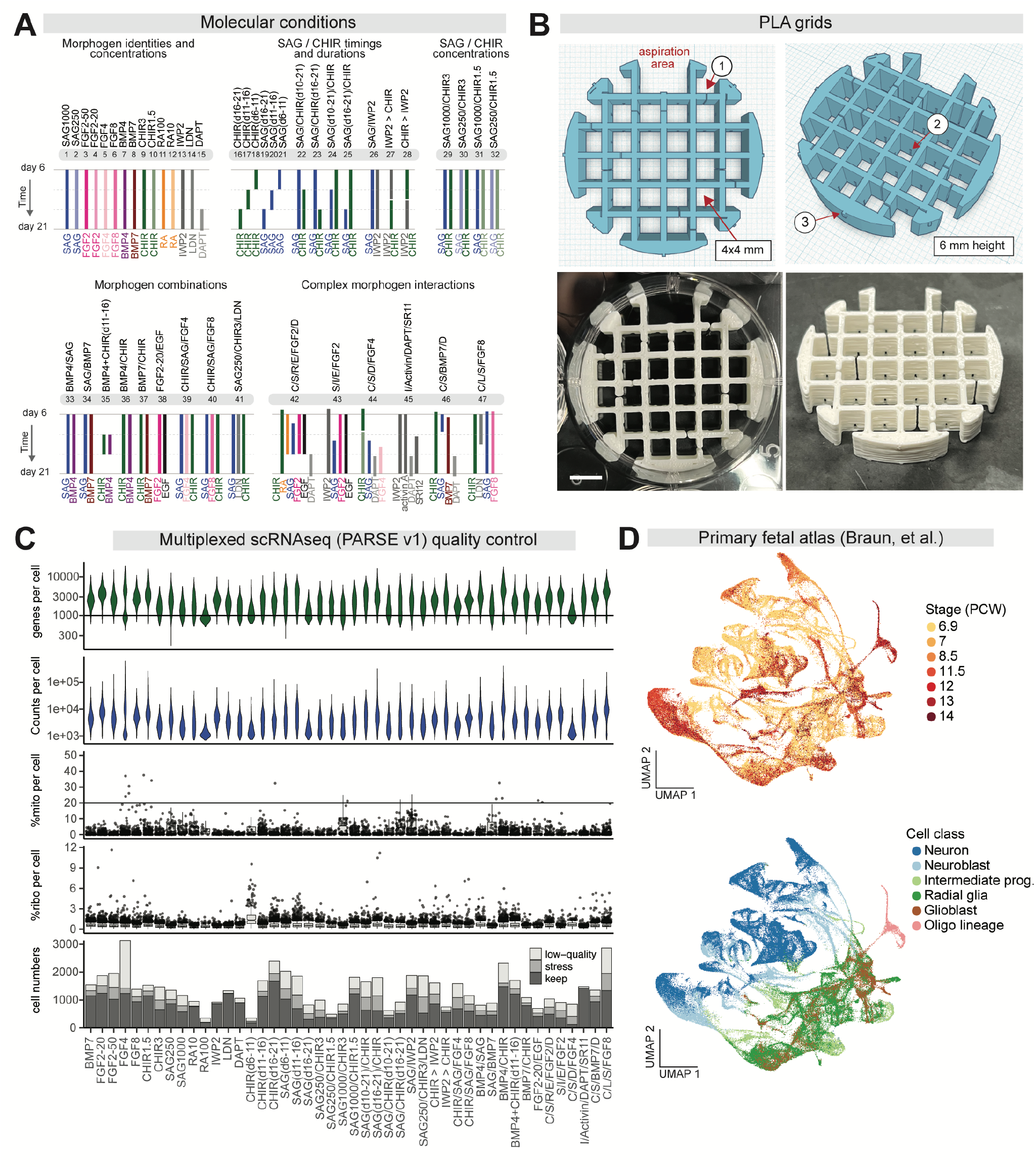
Schematic of organoid screen and single-cell RNA sequencing data quality. **(A)** Schematic of timing and dosing of each molecular condition (n = 47). See Supplementary Table 1. One condition (7, BMP4) resulted in organoids that progressively decreased in size and eventually disintegrated, likely due to differentiation into neural crest cells that leave the organoid; thus, n= 46 conditions were used for sequencing. **(B)** Top: design schematic of three-dimensional (3D) bio-fabricated polylactic acid (PLA) polymer grids for individual organoid differentiations. Bottom: representative images of printed grids inside culture well (left). 1. Open spaces to allow for compression when placed in a 6-well. 2. Diffusion ports. 3. Outer wings that wedge the grid into the 6 well plate to keep it in place. Scale bar, 4mm. **(C)** scRNA-seq quality metrics showing the distribution of the number of counts, number of genes, mitochondrial (mito) gene fraction, and ribosomal (ribo) gene fraction per cell in each sample. Mitochondrial and ribosomal gene fraction plotted as boxplots (horizontal line denotes median; lower and upper hinges correspond to the first and third quartiles; whiskers extend 1.5 times the interquartile range with outliers shown outside this range). Lines denote cell quality thresholds. Bottom: number of cells per sample meeting quality-thresholds and filtered for putative stressed cells (keep). **(D)** Same integrated UMAP as shown in Figure 1D, colored by primary human brain reference annotations. Top: sample age of each cell; PCW, post-conception week. Bottom: cell class annotations.

**Supplementary Figure 2.**
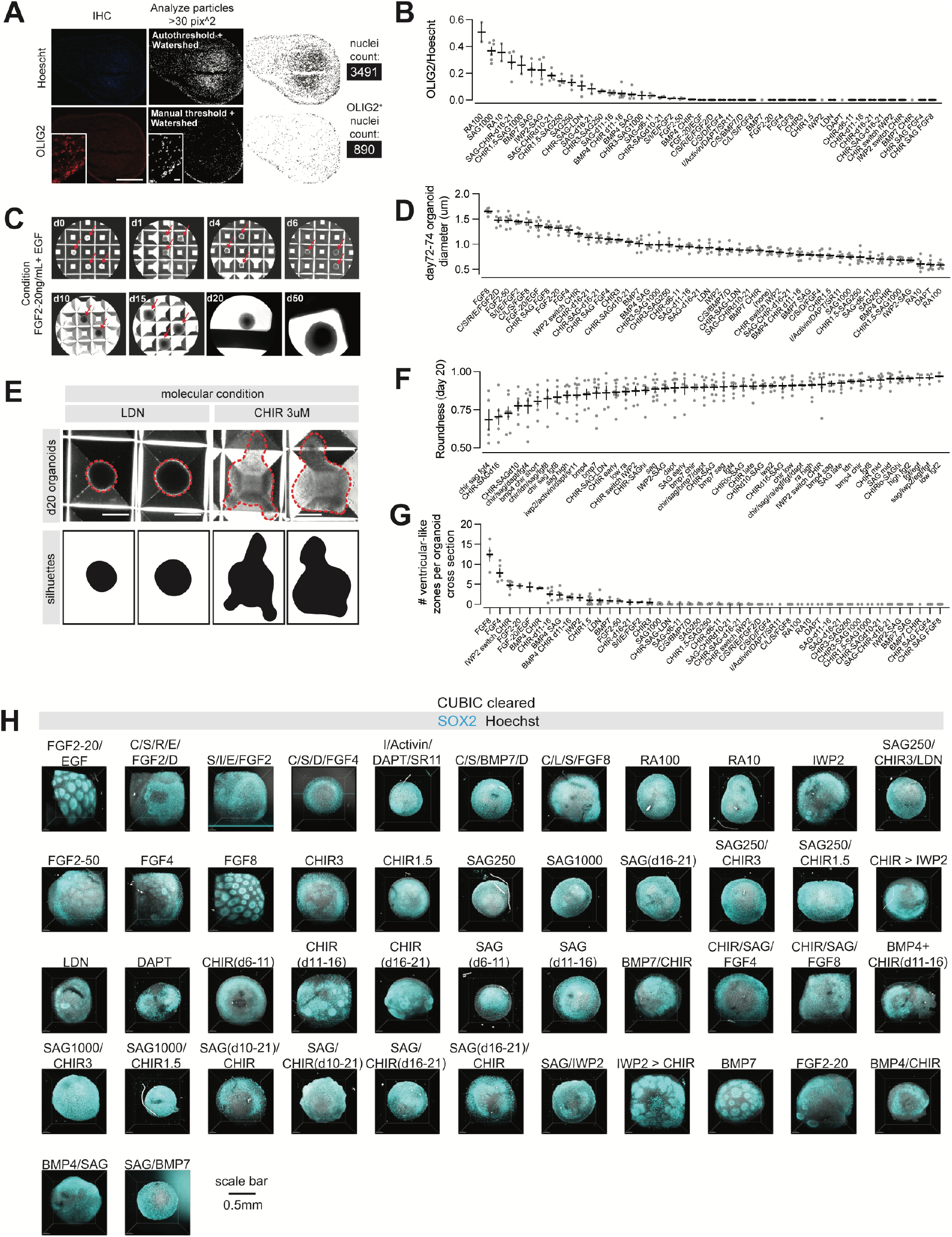
Cytoarchitectural characterization of organoid conditions. **(A)** Representative IHC image and analysis workflow for quantification of OLIG2 cell density. Scale bars, 0.2 mm, 25 nm. **(B)** Proportion of OLIG2^+^ nuclei at differentiation day 72–74 (determined by co-labeling with Hoechst) for each organoid condition (n= 1–7 organoids per condition). Data are presented as mean ± SEM. **(C)** Representative bright field images obtained over differentiation time. Scale bar, 0.8mm. **(D)** Diameter of each organoid (n= 1–7 organoids per condition). Data are presented as mean ± SEM. **(E)** Representative bright field images to illustrate morphological analysis workflow. Scale bar, 0.4 mm. **(F)** Roundness metric (ImageJ) of each organoid at differentiation day 20 (n= 1–7 organoids per condition). Data are presented as mean ± SEM. **(G)** Number of ventricular-like zones (methods) for each organoid (n= 1–7 organoids per condition). Data are presented as mean ± SEM. **(H)** CUBIC cleared images of SOX2 stained organoids for each condition at differentiation day 72–74. Scale bar, 0.5 mm.

**Supplementary Figure 3.**
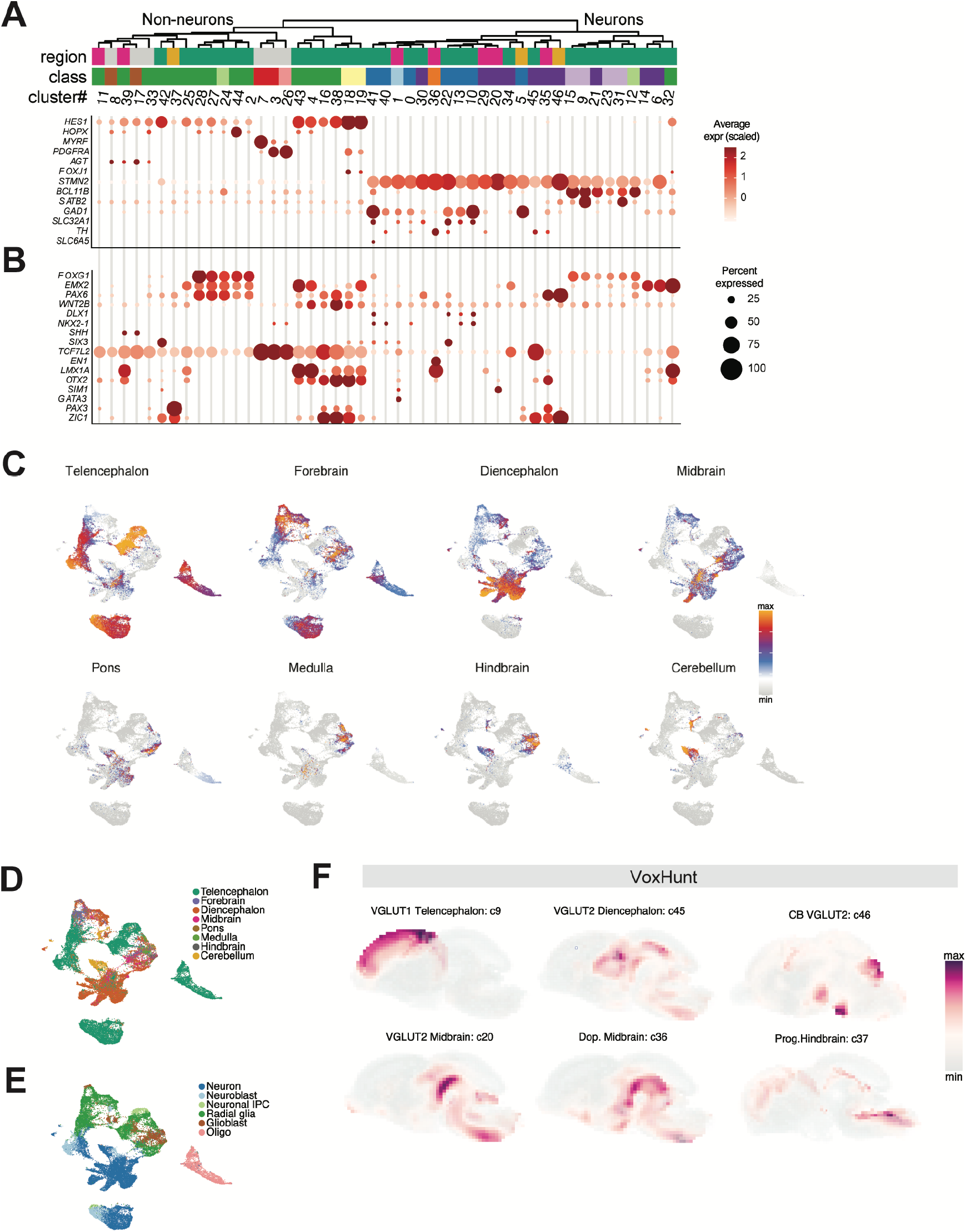
Single-cell RNA sequencing marker genes and cluster annotation. **(A)** Top: hierarchical clustering of scRNA-seq organoid clusters with brain region and cell class annotations. Bottom: Dot plot of example cell class marker gene expression. **(B)** Dot plot of example brain region marker gene expression. **(C)** UMAP visualization of developing human brain region classification scores for each organoid scRNA-seq cell using label transfer from primary human fetal reference scRNA-seq data (*28*). **(D)** UMAP visualization of predicted developing human brain region labels. **(E)** UMAP visualization of predicted developing human brain cell class labels. **(F)** VoxHunt (*34*) spatial brain mapping of the select organoid clusters onto developing mouse brain data from the Allen Brain Institute.

**Supplementary Figure 4.**
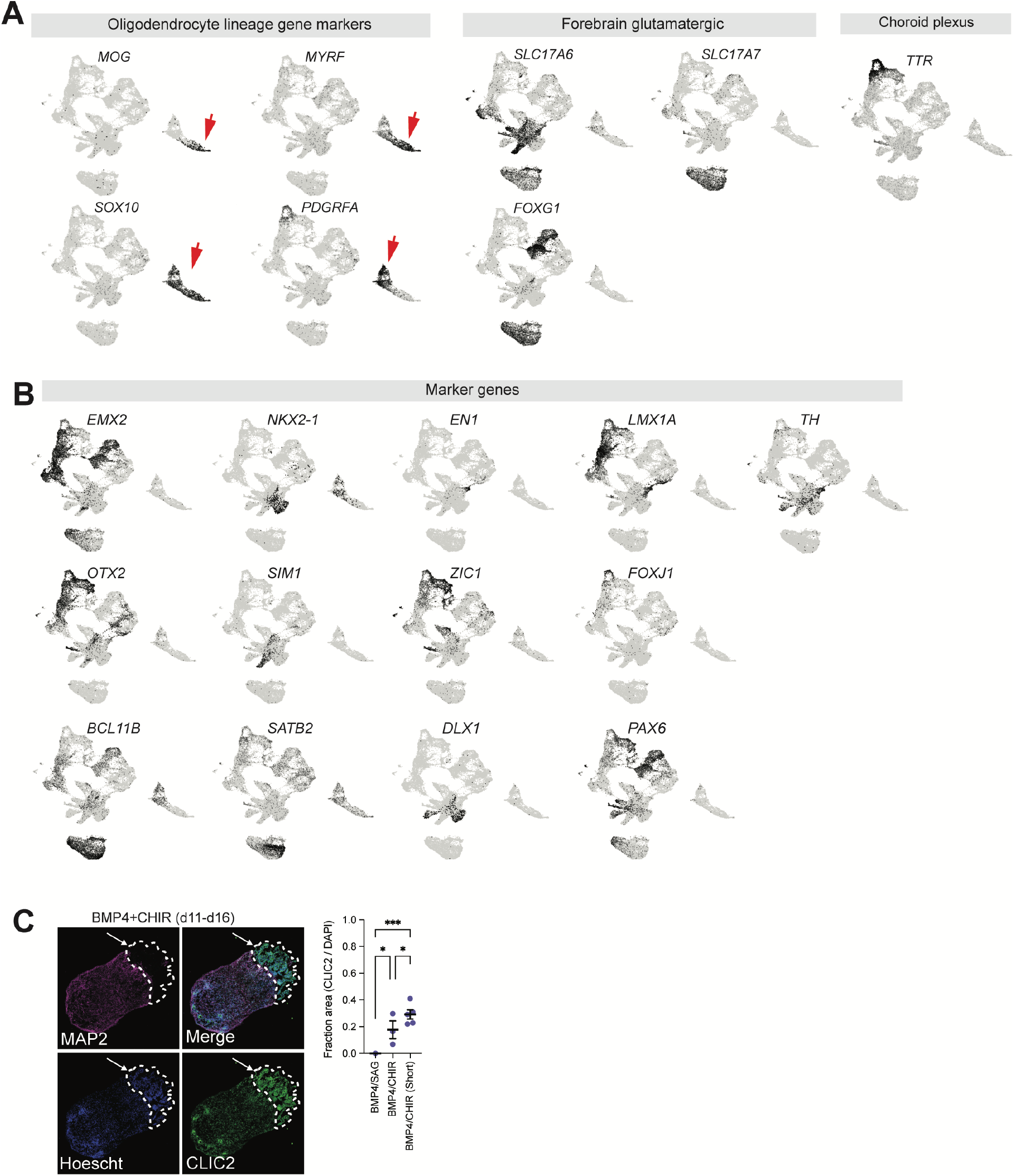
UMAP visualization of organoid scRNA-seq marker genes and histological validation. **(A)** UMAP visualization of expression for example cell class marker genes. **(B)** UMAP visualization of expression for example brain region marker genes. **(C)** Left: representative IHC images of choroid cell labeling from BMP4-CHIR (d11–16) condition at differentiation day 72. Arrow denotes CLIC2^+^ choroid plexus cells. Scale bar, 0.2 mm. Right: quantification of the proportion of CLIC2^+^ cells per condition (n= 3 images per organoid; two-sided t-test; P^*^ <0.05, ^**^<0.01, ^***^<0.001).

**Supplementary Figure 5.**
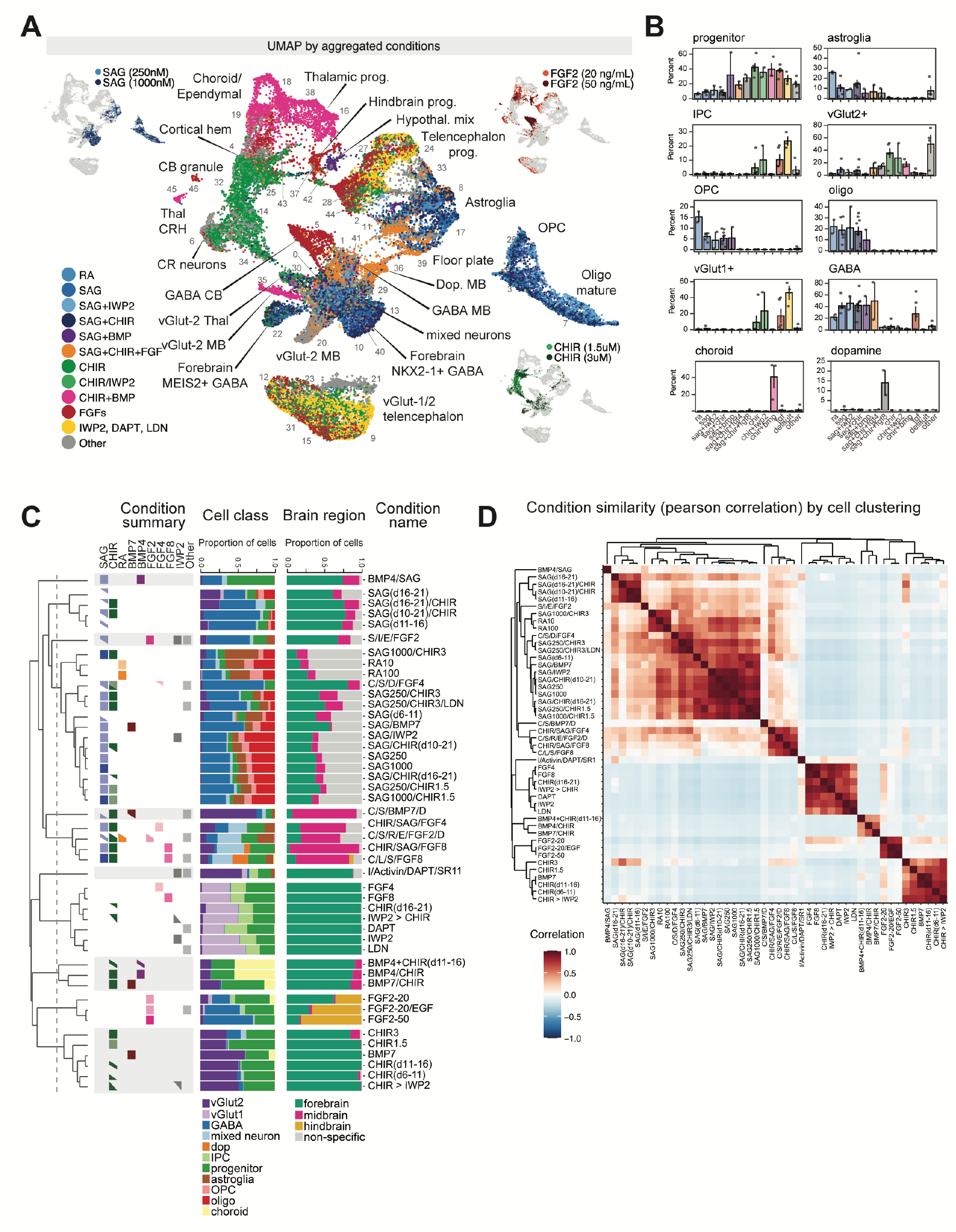
Molecular condition associations of organoid scRNA-seq clusters. **(A)** UMAP visualization of aggregated molecular condition information. Insets highlight specific condition doses for a given molecule. **(B)** Bar plot of the percentage representation of the indicated cell class in each aggregated molecular condition. Data points show individual organoid conditions. Bar plots denote mean ± SEM. **(C)** Left: hierarchical clustering of the cell cluster proportions across conditions. Select molecules are annotated for each condition. Middle: proportion of annotated cell class information per organoid condition. Right: proportion of annotated brain region information per organoid condition. **(D)** Heatmap of Pearson correlation matrix calculated from the cell cluster proportion representation of each condition.

**Supplementary Figure 6.**
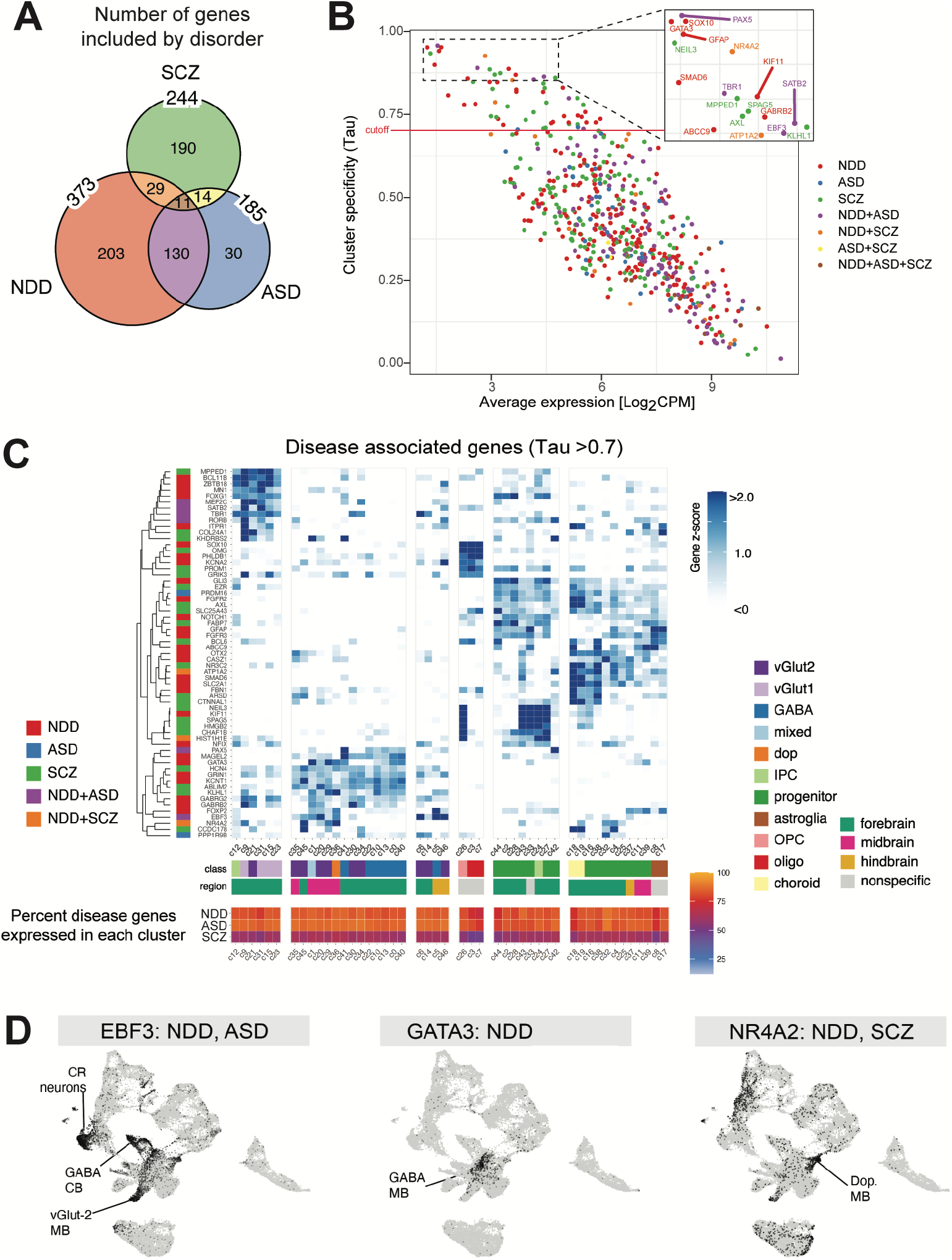
Neurodevelopmental disorder disease gene analysis. **(A)** Venn diagram of disease gene sets. **(B)** Scatter plot of average cluster expression (counts per million (CPM), log_2_ transformed) versus cluster specificity (Tau score, methods). Inset highlights high cluster specific disease genes. **(C)** Left: hierarchical clustering and gene set annotation of shown disease genes. Right: heatmap z-score normalized organoid cluster expression of select disease genes (Tau > 0.7). Bottom: organoid scRNA-seq cell class and region annotations. Heatmap of disease gene set percent expressed (defined as the non-zero expression in at least 10% of cells in a given cluster). **(D)** UMAP visualization of organoid scRNA-seq gene expression for select disease genes.

**Supplementary Figure 7.**
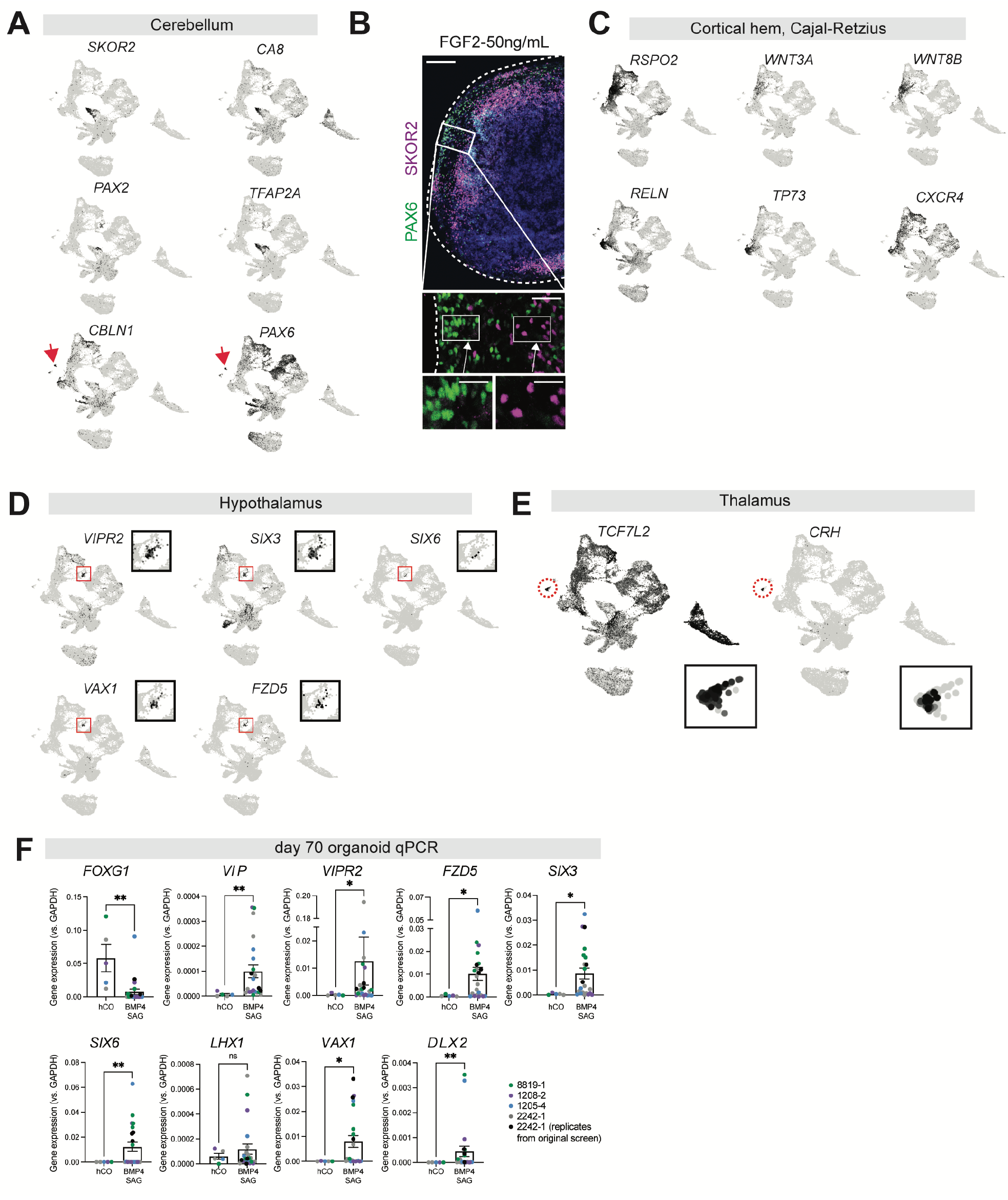
Generation of specific brain regions. **(A)** UMAP visualization of gene expression for selected cerebellar cell markers. Representative IHC images of cerebellar cell markers from the indicated condition. Scale bars, 0.1 mm (top), 100 nm (middle), 20 nm (bottom). UMAP visualization of gene expression for selected markers. **(D)** UMAP visualization of gene expression for selected hypothalamus markers. **(E)** UMAP visualization of gene expression for selected thalamus markers. **(F)** RT-qPCR of indicated hypothalamus markers in human cortical organoids (hCO) compared to BMP4-SAG condition across 4 hPS cell lines (n= 3 organoid per condition; two-sided t-test; P^*^ <0.05, ^**^<0.01, ^***^<0.001).

**Supplementary Figure 8.**
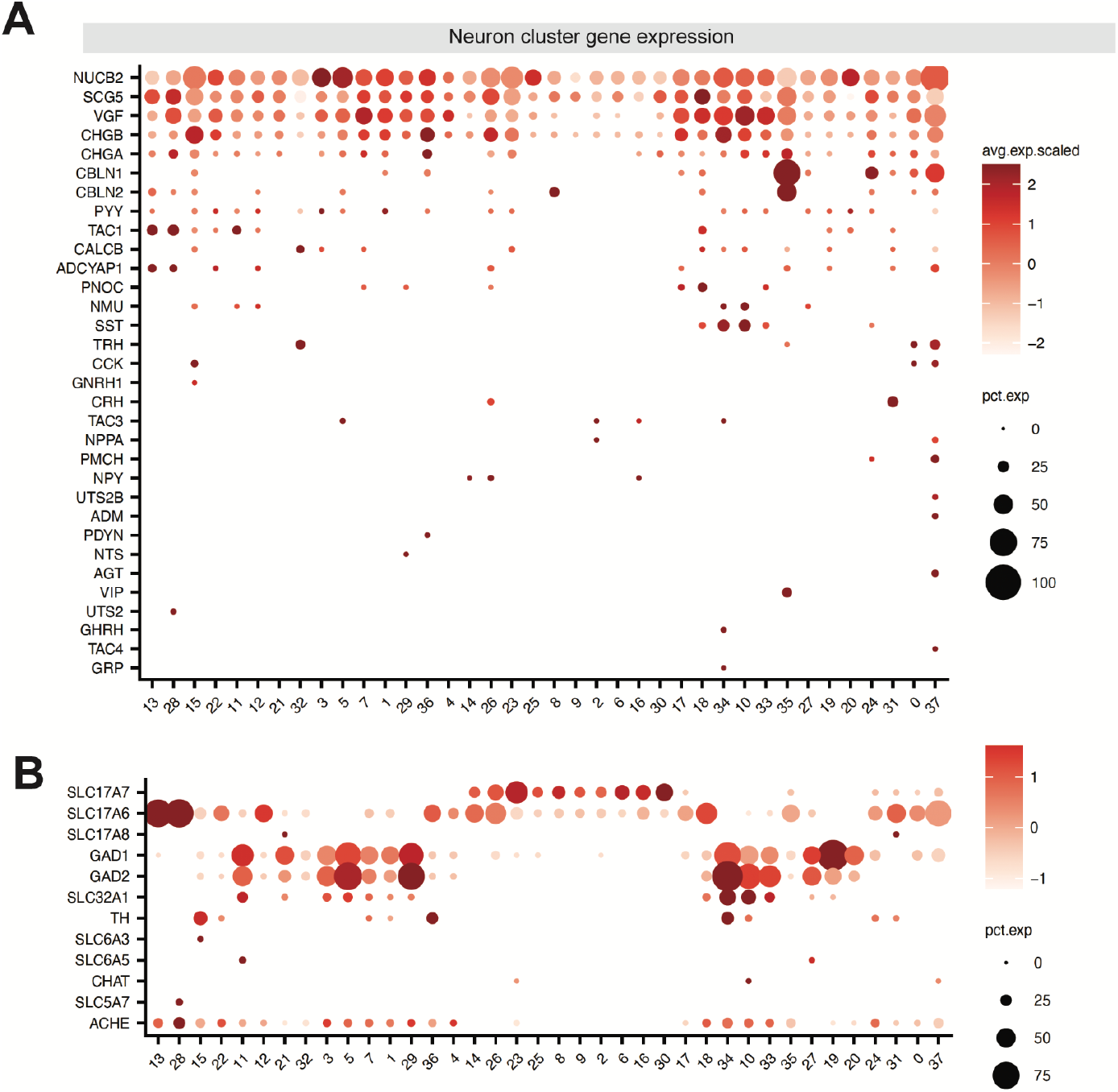
Expression of neuropeptide and neurotransmitter associated genes across organoid neuronal subclusters. **(A)** Dot plot visualization of neuropeptide associated genes ordered by the number of clusters with non-zero expression. Human neuropeptide genes were obtained from (*57*) and plotted if expressed in at least 10% of cells in at least one neuronal subcluster. **(B)** Dot plot visualization of neurotransmitter associated genes.

**Supplementary Figure 9.**
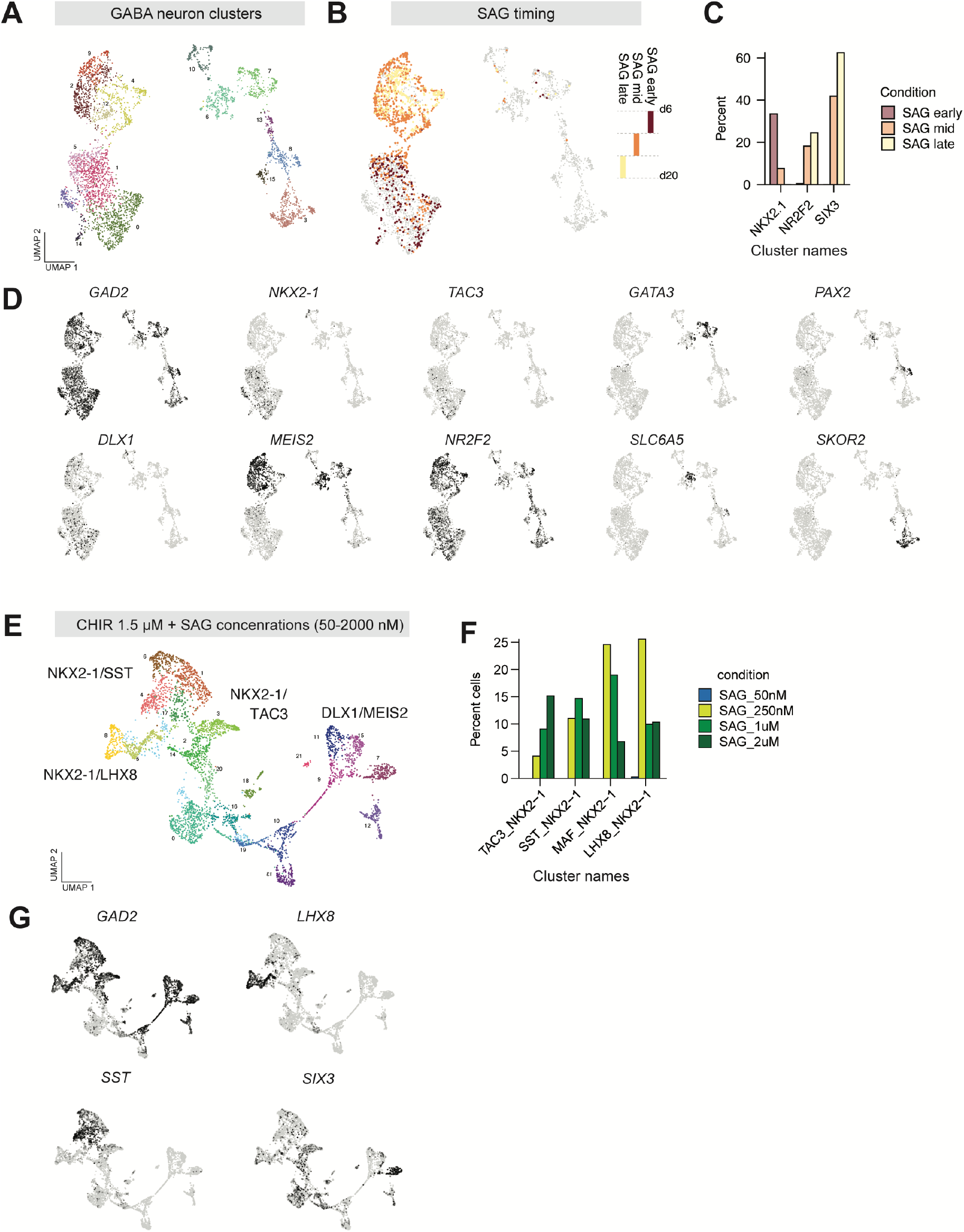
GABAergic neuron scRNA-seq in organoids and primary human fetal striatum. **(A)** UMAP visualization of GABAergic neuron subclusters from initial multiplex organoid screen. **(B)** UMAP visualization SAG timing condition information of GABAergic neuron cells from initial multiplex organoid screen. **(C)** Bar plot of percentage of cells from each SAG timing condition in the annotated cell groups. **(D)** UMAP visualization of gene expression from selected markers across GABAergic neuron cells from initial multiplex organoid screen. **(E)** UMAP visualization of scRNA-seq DLX1^+^ clusters from secondary GABAergic neuron organoid screen. **(F)** Bar plot of percentage of cells from each SAG concentration condition in the annotated cell groups. **(G)** UMAP visualization of gene expression for selected markers from secondary GABAergic neuron organoid screen.

